# Metaproteogenomic profiling of chemosynthetic microbial biofilms reveals metabolic flexibility during colonization of a shallow-water gas vent

**DOI:** 10.1101/2020.10.15.340729

**Authors:** Sushmita Patwardhan, Francesco Smedile, Donato Giovannelli, Costantino Vetriani

## Abstract

Tor Caldara is a shallow-water gas vent located in the Mediterranean Sea, with active venting of CO_2_, H_2_S. At Tor Caldara, filamentous microbial biofilms, mainly composed of *Epsilon*- and *Gammaproteobacteria*, grow on substrates exposed to the gas venting. In this study, we took a metaproteogenomic approach to identify the metabolic potential and *in situ* expression of central metabolic pathways at two stages of biofilm maturation. Our findings indicate that inorganic reduced sulfur species are the main electron donors and CO_2_ the main carbon source for the filamentous biofilms, which conserve energy by oxygen and nitrate respiration, fix dinitrogen gas and detoxify heavy metals. Three metagenome-assembled genomes (MAGs), representative of key members in the biofilm community, were also recovered. Metaproteomic data show that metabolically active chemoautotrophic sulfide-oxidizing members of the *Epsilonproteobacteria* dominated the young microbial biofilms, while *Gammaproteobacteria* become prevalent in the established community. The co-expression of different pathways for sulfide oxidation by these two classes of bacteria suggests exposure to different sulfide concentrations within the biofilms, as well as fine-tuned adaptations of the enzymatic complexes. Taken together, our findings demonstrate a shift in the taxonomic composition and associated metabolic activity of these biofilms in the course of the colonization process.

## Introduction

Marine gas vents transport volatile elements and compounds from the geosphere to the hydrosphere by seepage through sediments and bedrocks (Suess, 2014). At these sites, geological processes, biogeochemical reactions and the activity of microorganisms act together to alter the composition of volatile products. The release of such reduced volatiles into the oxygenated water column generates a redox disequilibrium that can be harnessed by benthic prokaryotic communities to convert chemical energy into ATP. Hydrothermal and gas vents occur both in shallow water (depth < 200 meters) and in the deep-sea (depth > 200 meters; Tarasov et al., 2005). Shallow-water hydrothermal and gas vents are associated with submarine volcanoes, arc and back-arc volcanoes and occur globally in the proximity of active plate margins and intraplate hotspots (reviewed in Price and Giovannelli, 2017).

The subduction of the African plate below Europe has resulted in the formation of the Mediterranean Ridge and deep subduction basins as well as active volcanic arcs in the Tyrrhenian and Aegean Seas (Dando *et al*., 1999). The terrestrial volcanic systems in these areas have been well-studied, both because of the long history of devastation caused by their eruptions and because of their geothermal potential. The submarine parts of the system have received relatively little attention until the last decade, and are still poorly studied compared to some of the mid-ocean ridge systems (Price and Giovannelli, 2017). Volcanic arc hydrothermal systems release large volumes of volatiles (Barry *et al*., 2019), because of the both degassing of the subducted slab and the mantle, and the decomposition of carbonates in the overlying marine sediments (Fisher, 2008). Since most of the known venting in the Mediterranean is from shallow vents, the majority of the outlets are of the gasohydrothermal type, emitting large volumes of carbon dioxide.

The microbiology of the shallow-water vent systems of the Aegean and Tyrrhenian basins has been mainly investigated in sediments (Giovannelli et al., 2012; Gugliandolo and Maugeri, 1993; Kerfahi et al.; Maugeri et al., 2013; Price et al., 2013; Sievert et al., 1999, 2000). However, besides sediment communities, substrate-attached chemosynthetic microbial biofilms are widespread and, in their role of primary producers, relevant in these ecosystems (Kalanetra *et al*., 2004; Reigstad *et al*., 2011; Miranda *et al*., 2016). Here, we investigated two types of biofilm communities at a shallow-water gas vent located at Tor Caldara, Italy.

Tor Caldara is a natural reserve located near the town of Anzio, about 60 km south of Rome. Active venting of gases, including CO_2_ originating from a deep magma source, occurs at Tor Caldara in association with the quiescent volcanic complex of Colli Albani, located around 40 km north-east of the venting site (Carapezza and Tarchini, 2007). The absence of a thermal anomaly at Tor Caldara is attributed to the infiltration of cold meteoric water (Carapezza *et al*., 2012). In a previous study, we investigated the chemical composition of the gases at Tor Caldara and the taxonomic diversity of the associated filamentous biofilm communities (Patwardhan *et al*., 2018). The gases at Tor Caldara are mainly composed of CO_2_ (avg. of 76.67 mol%) and H_2_S (avg. of 23.13 mol%) with minor contribution of CH_4_ (avg. of 0.18 mol%), CO (avg. of 0.0080 mol%) and H_2_ (avg. of 0.00072 mol%). Young and established filamentous biofilms - the former collected on glass slides deployed at the venting site at Tor Caldara and the latter attached to the native rocky substrates - were dominated by members of the classes *Epsilonproteobacteria* and *Gammaproteobacteria*, respectively, albeit in different proportions. On average, sulfur-oxidizing *Epsilonproteobacteria* of the genus *Sulfurovum* accounted for 57.6% of the active young biofilms, while sulfur-oxidizing *Gammaproteobacteria* of the genus *Thiomicrospira* as well as sequences related to the *Thiothrix* CF-26 group constituted more than 60% of the active established biofilm community (Patwardhan *et al*., 2018). This observed transition from *Epsilon-* to *Gammaproteobacteria* during the maturation of the biofilms revealed an ecological succession between the young and established communities. Previous studies of sulfidic environments, including caves, shallow-water and deep-sea hydrothermal vents, reported a spatial segregation between these two classes of bacteria and correlated this distribution pattern with geochemical data, leading to the hypothesis that *Epsiloproteobacteria* can cope with higher concentrations of hydrogen sulfide than *Gammaproteobacteria* (Engel et al., 2004; Giovannelli et al., 2013; Macalady et al., 2008; Meier et al., 2017; Miranda et al., 2016; O’Brien et al., 2015; Reigstad et al., 2011). Based on these studies, and on the observed succession observed at Tor Caldara, we tested the hypothesis that *Epsilon-* and *Gammaproteobacteria* are adapted to different concentrations of sulfide. Sulfide gradient experiments with laboratory strains showed that, conservatively, *Epsilonproteobacteria* can tolerate sulfide concentrations up to 20 times higher than *Gammaproteobacteria*, possibly facilitating their role as pioneer colonizers of sulfidic environments (Patwardhan *et al*., 2018).

To conserve energy, prokaryotes oxidize reduced inorganic sulfur compounds via a number of different pathways. The Sox pathway is widespread and occurs in both anaerobic as well as aerobic phototrophic and chemolithoautotrophic microorganisms. The well-characterized Sox enzyme complex in the alphaproteobacterium, *Paracoccus pantotrophus*, has four periplasmic proteins: SoxYZ, SoxXA, SoxB and SoxCD. These proteins work in concert to oxidize thiosulfate, hydrogen sulfide, sulfur as well as sulfite all the way to sulfate. In some sulfur-oxidizing bacteria (SOB) such as *Rhodobacter capsulatus*, sulfide is oxidized to sulfur by the membrane bound sulfide quinone oxidoreductase (SQR) enzyme. Additionally, in few SOBs, sulfide is also oxidized to sulfur by the periplasmic sulfide dehydrogenase (FccAB) (Visser *et al*., 1997; Mußmann *et al*., 2007)

In this study, we used an integrated metagenomic/metaproteomic approach to compare the metabolic potential and the *in situ* expression of central metabolic pathways of the established and young filaments at Tor Caldara. We identified and quantified key genes and proteins involved in the carbon, sulfur and nitrogen cycles and compared them between the young and established biofilms to establish how different metabolic pathways were expressed in the two communities. We also reconstructed three metagenome-assembled genomes (MAGs) belonging to the classes *Epsilonproteobacteria* and *Gammaproteobacteria*.

## Materials and Methods

### Study Site

Tor Caldara is a natural reserve located approximately 60 km south of Rome and 40 km south-west of Colli Albani, Italy (Patwardhan *et al*., 2018). The study site (41°29′ 9″ N 12°35′ 23″ E) is a coastal submarine gas vent located at a depth of approximately 3 meters. The site is characterized by vigorous venting of gases that escape from the sandy seabed. The sediment in the venting area is distinctly darker in color than the control sediment because of possible sulfide deposition. Conspicuous growth of white microbial filaments is seen on the rocks directly exposed to the gas venting.

### Sample Collection

Sampling of filaments was performed as described previously (Patwardhan *et al*., 2018). Briefly, established filaments (EF from here on) growing on rocks exposed to the gas venting were collected by a SCUBA diver in August 2016. To study the early stages of colonization, young biofilms (YF from here on) were collected on sterile glass slides mounted on an aluminum rod and exposed to the venting area for four days. Biofilms samples were then stored in RNA Later at −80°C for further nucleic acid and protein extraction. Representative samples of the two communities (one EF and one YF) were further used for downstream applications.

### Nucleic acid extraction, metagenomic library preparation and analysis

DNA was extracted from YF and EF biofilm biomass stored in RNA Later following a phenol:chloroform extraction protocol. Briefly, 0.8 g of sample were added to 850 μl of extraction buffer (50 mM Tris-HCl, 20 mM EDTA, 100 mM NaCl; pH 8.0) supplemented with100 μl of lysozyme (100 mg/ml) and incubated at 37°C for 30 min. This mix was then supplemented with 5 μl of proteinase K (20 mg/ml), incubated at 37 °C for 30 min, and subsequently supplemented with 50 μl SDS (20%) and further incubated at 65 °C for 1 hr. Nucleic acids were extracted by performing a series of phenol:chloroform:isoamylalcohol (25:24:1) and chloroform:isoamyl alcohol (24:1) extractions. Multiple samples were co-extracted to reduce potential bias. The final supernatant was precipitated in 3 M sodium-acetate and isopropanol, washed twice with 70% ice-cold ethanol and re-suspended in ultra-pure sterile water. The library was prepared at Molecular Research LP (Shallowater, TX), using the Nextera DNA Sample preparation kit (Illumina) following the manufacturer’s user guide. The initial concentration of DNA was evaluated using the Qubit^®^ dsDNA HS Assay Kit (Life Technologies). Initial library preparation resulted in very large size library (>2500 bp), therefore, library preparation was repeated after DNA inhibitor removal using the DNEasy PowerClean Pro Cleanup Kit (Qiagen). 50 ng DNA was used to prepare the library. The sample underwent the simultaneous fragmentation and addition of adapter sequences. These adapters are utilized during a limited-cycle (5 cycles) PCR in which unique indices were added to the sample. Following the library preparation, the final concentration of the library was measured using the Qubit^®^ dsDNA HS Assay Kit (Life Technologies), and the average library size was determined using the Agilent 2100 Bioanalyzer (Agilent Technologies). The library was diluted (to 10 pM) and sequenced paired end for 500 cycles using the HiSeq platform (Illumina). Following the removal of the adapters, at the sequencing facility, the sequences were quality checked using FastQC v.0.11.5 (Andrews S., 2010. FastQC: a quality control tool for high throughput sequence data: http://www.bioinformatics.babraham.ac.uk/projects/fastqc/). A total of 6,698,798 and 10,125,188 paired end sequences were used for downstream analysis for the EF and YF samples respectively. Sequences encoding rRNA genes were extracted using Metaxa2 v2.1 using default parameters (Bengtsson-Palme Johan *et al*., 2015) and annotated using Silva ngs v.1.3.9 (Quast *et al*., 2013). The paired end sequences were then assembled using Megahit v1.1.3 (Li *et al*., 2015) with default parameters, and assembly was checked using QUAST v3.4 (Gurevich *et al*., 2013). The EF assembly had a total of 99,137 contigs with the largest contig being 212,020 bp long, whereas the YF assembly had a total of 87,836 contigs with the largest contig being 261,286 bp long. Assembled metagenomes were submitted to DOE Joint Genome Institute’s Integrated Microbial Genome Metagenomic Expert Review (IMG/ER) pipeline for Open Reading Frame (ORF) identification as well as functional and taxonomic annotation (Markowitz *et al*., 2012). The EF and YF metagenomic assemblies had a total of 308,874 and 362,063 sequences, respectively. The sequences of both metagenomes had COG (51%), KO (41%) and KEGG (25%) annotations. Post annotation, using default parameters for bowtie2 v.2.3.3.1, the reads were mapped back to the assembly (Langmead *et al*., 2009). Using HTSeq v0.9.1, the number of reads mapping to each gene was counted (Anders *et al*., 2015). <u>Custom scripts</u> were used to normalize absolute read counts mapped to each gene for varying gene lengths and difference in sequencing depth between samples. These normalized gene abundances were expressed as TPM (Transcripts Per Million; (Wagner *et al*., 2012). Sequences are available through the NCBI Short Read Archive database with accession number PRJNA498803.

### Protein extraction and metaproteomic analysis

Total proteins were extracted from the YF and EF biofilms by addition of Laemmli buffer, followed by sonication and incubation at 95 °C. Proteins were then precipitated by addition of urea, purified and digested with trypsin on a SDS-PAGE gel, and then label-free samples were run on the LC-MSMS. The resulting tandem mass spectra were searched against the predicted peptide sequences encoded by the combined metagenomes of the two filament samples, using the open-source software X!Tandem (Craig and Beavis, 2004). A total of 138,387 and 97,607 peptides were obtained in the metaproteomes of EF and YF, respectively. The proteins identified via X!Tandem were functionally and taxonomically annotated using the corresponding annotated metagenomes. Protein counts were normalized to the total proteins obtained from each sample. Normalized counts for proteins of interest were manually compared between the two samples. These normalized counts were expressed as a percent of total proteins observed in each sample. Raw data was uploaded to the ProteomeXchange database.

### Whole genome reconstruction from metagenomes

Quality checked reads from both samples were co-assembled using Megahit v1.1.3 with default parameters and QUAST v3.4 was used to check the quality of the co-assembly. The co-assembled contigs from the two metagenomes were then binned using MaxBin2.0 (Wu *et al*., 2014) to recover individual genomes. Completeness, taxonomic affiliation and contamination of recovered bins was checked using CheckM v1.0.7 (Parks *et al*., 2015).

### Phylogenetic Analyses

Sulfide quinone oxidoreductase (SQR) amino acid sequences obtained in this study and from GenBank were aligned with ClustalX v 2.0 (Thompson *et al*, 1997) and manually adjusted using Seaview (Galtier *et al*, 1996). Phylogenetic distances were calculated using the LG amino acid replacement matrix (Le and Gascuel, 2008) and the maximum likelihood method was used to evaluate tree topologies in PhyML 3.0 (Guindon *et al*., 2010). Branch support was estimated using the approximate likelihood ratio test (aLRT; (Anisimova and Gascuel, 2006) and by bootstrap analysis with 1,000 resamplings. Both estimates gave comparable results.

## Results

### Microbial community composition of the established and young filaments

The bacterial diversity of EF and YF was evaluated following in-silico extraction of the 16S rRNA gene sequences from the two metagenomes. The EF biofilm was dominated by *Gammaproteobacteria* (54%), followed by *Epsilonproteobacteria* (29%) and small proportions of *Bacteroidia* (4%) and *Alphaproteobacteria* (3%; Fig.1a). Within the *Gammaproteobacteria*, the genus *Thiomicrospira* (40%; now reclassified as *Thiomicrorhabdus;* Boden et al., 2017) and uncultured members belonging to the *Thiotrichales* (11%) were dominant (Fig.1b). A reverse trend was observed in the YF biofilm, with *Epsilonproteobacteria* (67%) dominating the community, followed by *Gammaproteobacteria* (23%), *Bacteroidia* (2%) and *Alphaproteobacteria* (1%; Fig.1a). In the YF biofilm*, Sulfurovum* (66%) was the most abundant genus within the *Epsilonproteobacteria* (Fig.1b).

**Fig. 1.**
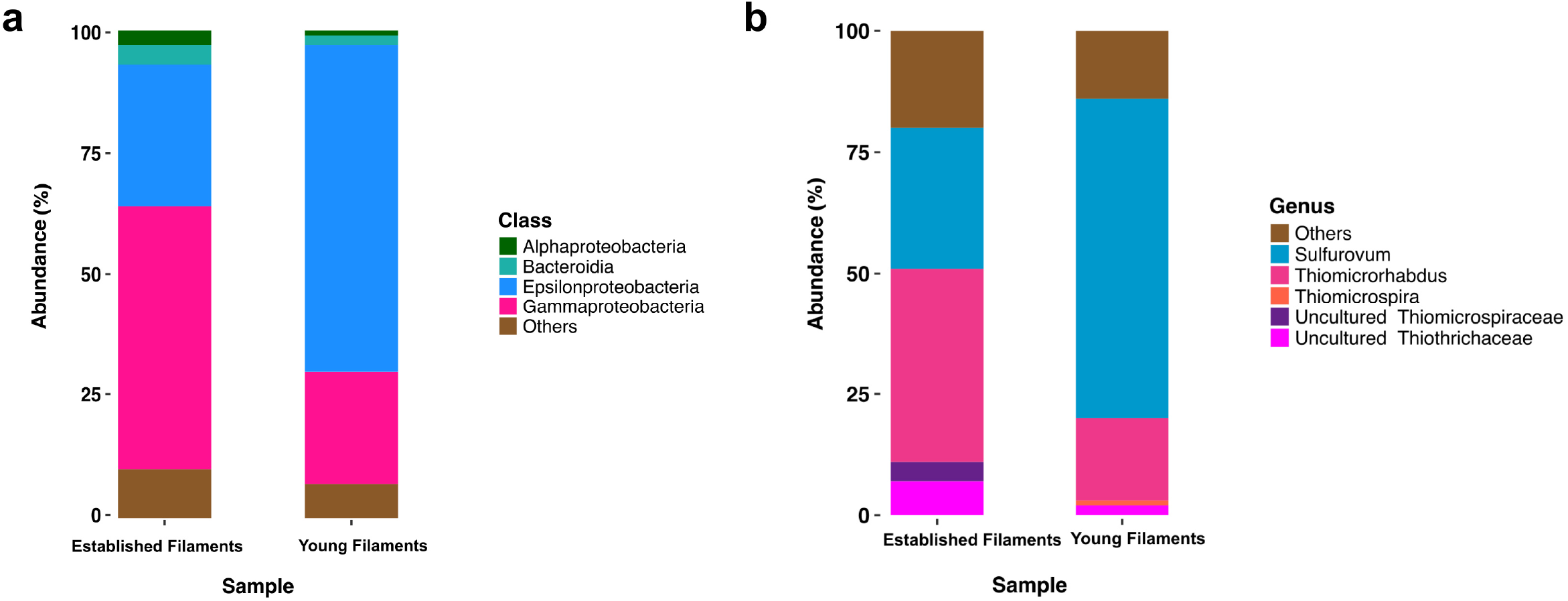
Class (a) and genus (b) level distribution of 16S rRNA gene and transcript sequences recovered from established and young filaments.

### Comparative metagenomics and metaproteomics of the established and young filaments

We compared the normalized abundances of key genes and proteins involved in central metabolic pathways, including carbon fixation, nitrogen, sulfur metabolism and (micro)aerobic respiration (Suppl. Tables 1 and 2). Since shallow-water vents are enriched in heavy metals, genes and proteins involved in their detoxification were also identified and compared (Suppl. Tables 1 and 2). In line with the 16S rRNA gene-inferred community composition, the majority of functional genes and proteins were affiliated with *Gamma-* and *Epsilonproteobacteria*, (Figs. 2, 3 and 4). The metagenomes provided a snapshot of the metabolic potential of the microbial communities, and were also used to annotate the metaproteomes, which revealed the *in situ* expression of central metabolic pathways of the communities. The metagenomes were also used to obtain metagenome-assembled genomes (MAGs) of the most abundant members of the filamentous biofilms.

**Fig. 2.**
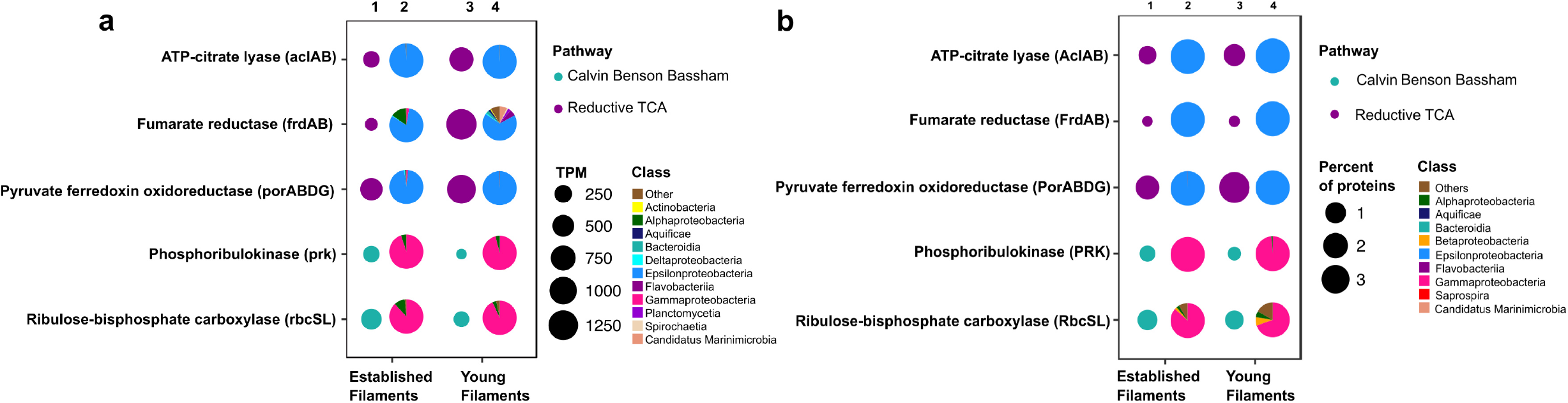
Profiles of genes and proteins for carbon fixation in established and young filamentous biofilms. (a) Transcripts per million (TPM; columns 1 and 3) and taxonomic affiliation of reads annotated to genes (columns 2 and 4); (b) Percentage (columns 1 and 3) and taxonomic affiliation of reads annotated to proteins (columns 2 and 4).

**Fig. 3.**
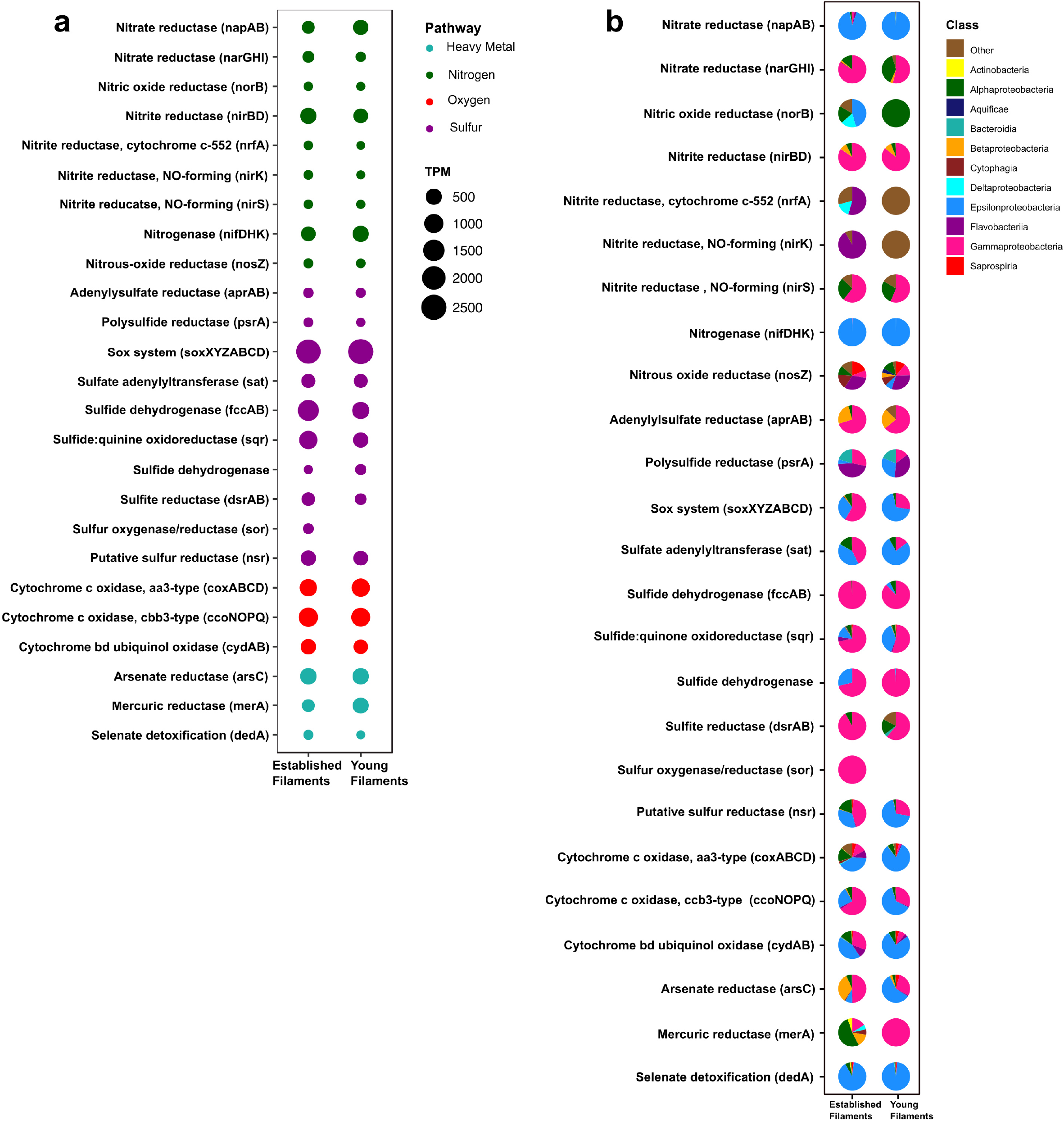
(a) Transcripts per million (TPM) and (b) taxonomic affiliation of reads annotated to key genes involved in nitrogen, sulfur, oxygen and heavy metal detoxification pathways for established and young filaments.

**Fig. 4.**
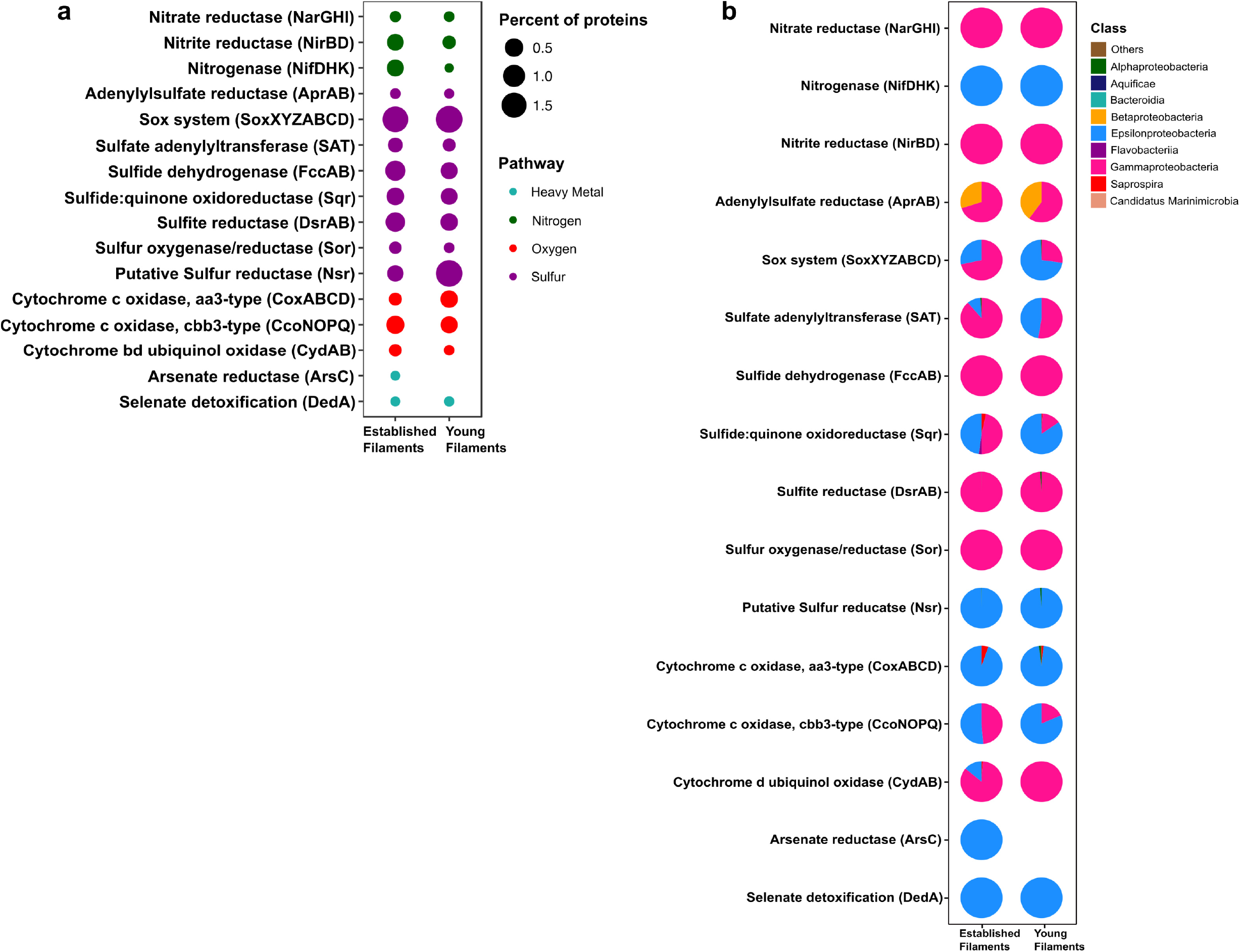
(a) Percentage and (b) taxonomic affiliation of key proteins involved in nitrogen, sulfur, oxygen and heavy metal detoxification pathways for established and young filaments.

#### Carbon fixation

The metagenome and the metaproteome of both filamentous communities included genes and proteins involved in the reductive tricarboxylic acid (rTCA) and Calvin-Benson-Bassham (CBB) cycles for carbon fixation (Fig. 2). The YF biofilm community had three times as many genes for the rTCA than the EF biofilm, while EF had twice as many genes for the CBB cycle than YF (Fig. 2a, columns 1 and 3, and Suppl. Table 1). While proteins of the rTCA cycle contributed only to 2.2% of the total proteins in EF, they contributed to 5% of the total in YF (Fig. 2b, columns 1 and 3, and Suppl. Table 2). Taxonomically, the rTCA proteins were all affiliated with *Sulfurovum* and *Sulfurimonas* within the *Epsilonproteobacteria* (Fig. 2b, columns 2 and 4). In contrast, proteins of the CBB cycle comprised 1.1% and 0.7% of the total proteins in EF and YF, respectively (Fig 2b, columns 1 and 3, and Suppl. Table 2), with most of them taxonomically identified as *Gammaproteobacteria* and specifically affiliated to the genera *Thiomicrospira, Thiothrix* and *Thioalkalivibrio* (Fig. 2b, columns 2 and 4). These bacteria are chemolithotrophs that derive energy from the oxidation of reduced sulfur species and are key players in marine geothermal environments.

#### Nitrogen and sulfur metabolism

The metagenomes of the two biofilm communities encoded for genes involved in nitrate reduction, denitrification, nitrogen fixation, sulfur oxidation/reduction, oxygen respiration and heavy metal detoxification (Fig. 3a). However, the abundance of some of these genes differed between the two biofilm communities. For instance, the *napAB* (encoding two components of the periplasmic nitrate reductase complex) and *nifDHK* (nitrogenase) genes, involved in nitrate reduction and nitrogen fixation, respectively, were found to be enriched in the YF biofilm (Fig. 3a and Suppl. Table 1) and were taxonomically affiliated with the *Epsilonproteobacteria* (Fig. 3b). The *narGHI* (membrane bound nitrate reductase) and *nirS* (nitrite reductase) genes, as well as those for sulfur oxidation, *fccAB* (sulfide dehydrogenase), and sulfate reduction/sulfide oxidation, *dsrAB* (dissimilatory bisulfite reductase) were enriched in EF (Fig. 3a and Suppl. Table 1). The *sor* gene (sulfur oxygenase/reductase) was found in EF but was absent in YF. Taxonomically, the *nar, nir* and *sor* genes were mostly affiliated with the *Gammaproteobacteria* (Fig. 3b). The *sqr* (sulfide quinone oxidoreductase) gene was also enriched in the EF biofilm community and predominantly affiliated with the *Gammaproteobacteria* (Figs. 3a and b). However, the proportion of *Epsilonproteobacteria*-affiliated *sqr* genes increased in the YF community (Fig. 3b).

In the metaproteome, the *Epsilonproteobacteria-affiliated* nitrogenase (Nif) accounted for 0.35% of the total proteins in EF as opposed to 0.001% in YF (Fig. 4a and Suppl. Table 2). The periplasmic nitrate reductase (Nap) was undetectable in both the samples, while both the membrane bound nitrate reductase (Nar) and the nitrite reductase (Nir), affiliated with the *Gammaproteobacteria*, were more abundant in EF (0.02% and 0.3%; Fig. 4a). Proteins involved in the dissimilatory sulfite reductase (Dsr) pathway for either elemental sulfur oxidation or sulfite reduction and the APS pathway for indirect sulfite oxidation (Apr, SAT) were twice as abundant in the EF biofilm compared to the YF one (Fig. 4a and Suppl. Table 2). Dsr and Apr in both metaproteomes were mostly affiliated with *Gammaproteobacteria*, while SAT was mostly affiliated with *Gammaproteobacteria* in the EF biofilm and with both *Gammaproteobacteria* and *Epsilonproteobacteria* in the YF community (Fig. 4b). The sulfide dehydrogenase (Fcc) was twice more abundant in the EF than in the YF biofilm (Fig. 4a) and it was affiliated with *Gammaproteobacteria* in both samples (Fig. 4b and Suppl. Table 2). In contrast, the abundance of the sulfide quinone oxidoreductase (SQR) was comparable between the two biofilms (Fig. 4a), but its taxonomic affiliation changed from comparably epsilon- and gammaproteobacterial in the EF biofilm to predominantly epsilonproteobacterial in the YF community (Fig. 4b and Suppl. Table 2). The Sor protein, involved in the disproportionation of elemental sulfur (Kletzin, 1989), was classified as *Thioalkalivibrio* spp. and was enriched in the EF biofilm (Fig. 4a and Suppl. Table 2). A putative NADPH-dependent sulfur reductase (NSR) affiliated with *Sulfurovum* spp. was six times more abundant in the total proteins of YF than EF (Fig. 4a and Suppl. Table 2). The *sox* genes and the Sox proteins were almost equally abundant in both the YF and EF biofilms (Figs. 3a and 4a); however, in the YF biofilm, both *sox* genes and Sox proteins were predominantly affiliated with the *Epsilonproteobacteria*, while in the EF biofilm they were mostly affiliated with the *Gammaproteobacteria* (Figs. 3b and 4b). We did not find any hydrogen oxidation-related genes or proteins in the metagenomes and metaproteomes of the two biofilm communities.

#### Oxygen reduction

Three cytochrome oxidase gene clusters (*coxABCD, ccoNOP* and *cydAB*) were abundant and comparable between the two metagenomes, indicating that oxygen respiration played an important role in both biofilm communities (Fig. 3a and Suppl. Table 1). In the YF biofilm, the majority of all three genes were taxonomically classified as *Epsilonproteobacteria*, while in the EF biofilm the taxonomic affiliation of *coxABCD* and *cydAB* revealed a higher diversity of the oxygen respiring community, except for *ccoNOP*, which was mostly affiliated with the *Gammaproteobacteria* (Fig. 3b). Compared to proteins involved in nitrate/nitrite respiration, the aa3-type and cbb3-type cytochromes, mediating oxygen respiration, were highly enriched (seven times as much) in YF (Fig. 4a and Suppl. Table 2). The majority of these proteins were classified as *Epsilonproteobacteria* (Fig. 4b) and were specifically affiliated with *Sulfurovum* spp., with the exception of cytochrome cbb3 in EF, which was equally affiliated with *Epsilon*- and *Gammaproteobacteria* (Fig. 4b). The cytochrome d (CydAB) was mostly affiliated with *Gammaproteobacteria* (Fig. 4b).

#### Heavy metal detoxification

Elevated concentrations of potentially toxic heavy metals such as mercury, arsenic, selenium etc. are present at hydrothermal vents, leading to heavy metal resistance in the resident microbiota (Rathgeber *et al*., 2002; Vetriani *et al*., 2005; Price *et al*., 2013). Genes involved in the detoxification of arsenate (*arsC*), mercury (*merA*) and selenate (*dedA*) were recovered from both metagenomes (Fig. 3a). The abundance of the *arsC* gene was comparable in both samples (Fig. 3a and Suppl. Table 1), while this gene was affiliated mostly with *Gammaproteobacteria* and *Epsilonproteobacteria* in the EF the YF, respectively (Fig. 3b). The *merA* gene (mercury reduction) was more abundant in the YF than the EF metagenome (Fig. 3a and Suppl. Table 1), and was affiliated with the *Gammaproteobacteria*, while the *dedA* gene (selenate detoxification) was more abundant in the EF and it was mostly affiliated with the *Epsilonproteobacteria* (Figs. 3a and b). In the metaproteome, only DedA, which has been shown to be involved in selenite resistance (Ledgham *et al*., 2005), was expressed in the YF biofilm, as opposed to both DedA and arsenate reductase (Ars) being expressed in EF (Fig. 4a and Suppl. Table 2). Both enzymes were classified as *Epsilonproteobacteria* (Fig. 4b). While present in both metagenomes, the mercuric reductase gene (*merA*) was not found in the metaproteomes of either biofilm communities (Figs. 3a and b).

### Metagenome-assembled genomes (MAGs)

Despite the fundamental role as primary producers of Gamma and Epsilonproteobacteria in marine geothermal habitats, there is a dearth of representative cultures and their genomes. A total of 31 MAG bins were obtained from the co-assembly of the two metagenomes. Out of these, three (TCMF1, TCMF2 and TCMF9) were ≥ 85% complete with contamination of ≤ 8% and complied with the proposed criteria that define medium to high quality draft MAGs (Bowers *et al*., 2017). The 16S rRNA genes were recovered from bins TCMF2 and TCMF9 (Table 1). The 16S rRNA gene of MAG TCMF2 showed >97% sequence similarity to 16S rRNA genes related to the *Thiotrix*-related CF-26 group recovered from microbial mats at shallow-water vents located on the South Tonga Arc (HQ153886) (Murdock *et al*. 2010) and at the Kuieshan Island (JQ611107). In our previous study of the Tor Caldara vents, dominant OTUs annotated as *Thiotrix*-related CF-26 from EF showed similar sequence similarity to the OTUs recovered from the same microbial mats, indicating that the TCMF2 MAG likely belongs to this group of uncultured *Thiotrichales*. Most of the genes involved in the CBB cycle, including RuBisCo (*rbcSL*), were present in MAG TCMF2. Further, the potential for thiosulfate oxidation was evidenced by the presence of *sox* genes, in addition to the *sqr* and *fccAB* genes involved in sulfide oxidation. This MAG also contained the genes for cbb3-type cytochrome oxidase and *nirBD*, involved in microaerobic and nitrite respiration, respectively (Table 1).

**Table 1.** Genes involved in main metabolic pathways recovered from MAGs.

| Pathway | Gene | MAG |  |  |
| --- | --- | --- | --- | --- |
|  |  | TCMF1<br>*Sulfurovum* | TCMF2<br>*Thiotrix*-related | TCMF9<br>Gammaproteobacteria |
|  | 16S rRNA | - | Yes | Yes |
| Calvin cycle | *rbc* | - | Yes | Yes |
|  | *prk* | - | - | Yes |
| Reductive TCA cycle | *kor* | Yes | - | - |
|  | *frd* | Yes | - | - |
| Sulfur oxidation | *sox* | Yes | Yes | Yes |
|  | *sqr* | Yes | Yes | Yes |
|  | *fcc* | - | Yes | Yes |
|  | *dsr* | - | - | Yes |
|  | *apr* | - | - | Yes |
|  | *sat* | - | - | Yes |
| Oxygen respiration | *cox* | Yes | - | Yes |
|  | *cco* | Yes | Yes | Yes |
|  | *cyd* | Yes | - | - |
| Nitrogen metabolism | *nir* | - | Yes | - |
|  | *nif* | Yes | - | - |
| Quorum sensing | *lux* | Yes | - | - |

MAG TCMF9 showed 93% 16S rRNA gene similarity to a gammaproteobacterial endosymbiont clone of the gutless oligochaete worm, *Olavius ilvae. O. ilvae* is known to harbor sulfur-oxidizing gammaproteobacterial endosymbionts that fix CO_2_ via the CBB cycle (Ruehland *et al*., 2008). This MAG contained most of the genes involved in the CBB cycle, including RuBisCo (*rbcSL*) and phosphoribulokinase (*prk*). A whole suite of genes required for thiosulfate and sulfide oxidation such as the *sox* system (except *soxC*), *sqr* and *fccAB* were present. This MAG also contained all three genes (*dsrAB, aprAB, sat*) involved in the reverse *dsr* pathway, indicating versatility for the oxidation of sulfur compounds. Three out of the four genes encoding the aa3-type cytochrome oxidase (*cox*) and all genes encoding the microaerobic cbb3-type cytochrome oxidase (*cco*) were present (Table 1). The majority of the genes involved in bacterial chemotaxis and flagellar motility were also present (data not shown).

Since the 16S rRNA gene was not recovered in MAG TCMF1, other single-copy ribosomal proteins were used to constrain its lineage. Both the 30S ribosomal protein S8 and S9 were greater than 90% identical to a *Sulfurovum* species. Additionally, the average amino acid identity (AAI) shared between TCMF1 and *Sulfurovum* strain NBC37 was 58%, which is well within the range that defines the genus boundary (Rodriguez-R and Konstantinidis, 2014). Based on this data, TCMF1 belonged to the *Sulfurovum* lineage within the *Epsilonproteobacteria*, which dominated the YF community (Fig. 1b). *Sulfurovum* spp. are sulfur/hydrogen oxidizers that fix CO_2_ using the rTCA cycle (Inagaki, 2004; Mino *et al*., 2014; Giovannelli *et al*., 2016). Out of the three diagnostic genes for the rTCA cycle, *kor* and *frd* were present. *Sox* and *sqr* genes were also present, while hydrogenases were absent. Interestingly, this MAG contained the *nifDHK* gene essential for nitrogen fixation, while the periplasmic nitrate reductase encoding gene, *napAB*, which is widespread in the *Epsilonproteobacteria* (Vetriani *et al*., 2014), was missing. Three out of the four genes encoding the aa3-type cytochrome oxidase (*cox*) and all genes encoding the micro-aerobic cbb3-type (*cco*) and bd cytochrome oxidase (*cyd*) were present, indicating the ability to respire oxygen within a broad range of concentrations. The *luxS* gene, involved in quorum sensing and conserved across all members of the *Epsilonproteobacteria* (Pérez-Rodríguez *et al*., 2015), was also present in TCMF1.

Given the presence of genes coding for *sqr* both in the metagenomes and in the recovered MAGs, we carried out phylogenetic analyses of representative SQR enzymes from the Tor Caldara biofilms and close relatives obtained from GenBank. This analysis placed the sequences into three discrete clusters separated from the *Chlorobium* lineage, from which SQR was originally characterized: one cluster was related to *Sulfurovum* spp., one was related to the *Thiotricales* (*Thiomicrospira* and *Thiothrix* spp) and a third cluster included two groups of sequences: one related to *Sulfurovum* spp. and one related to the *Thiotricales* (Fig. 5). In line with the taxonomic annotation derived from other genes, the SQR from MAG TCMF1 was related to *Sulfurovum* sequences, while the enzymes from MAGs TCMF2 and TCMF9 were related to *Thiotricales*-derived sequences (Fig. 5).

**Fig. 5.**
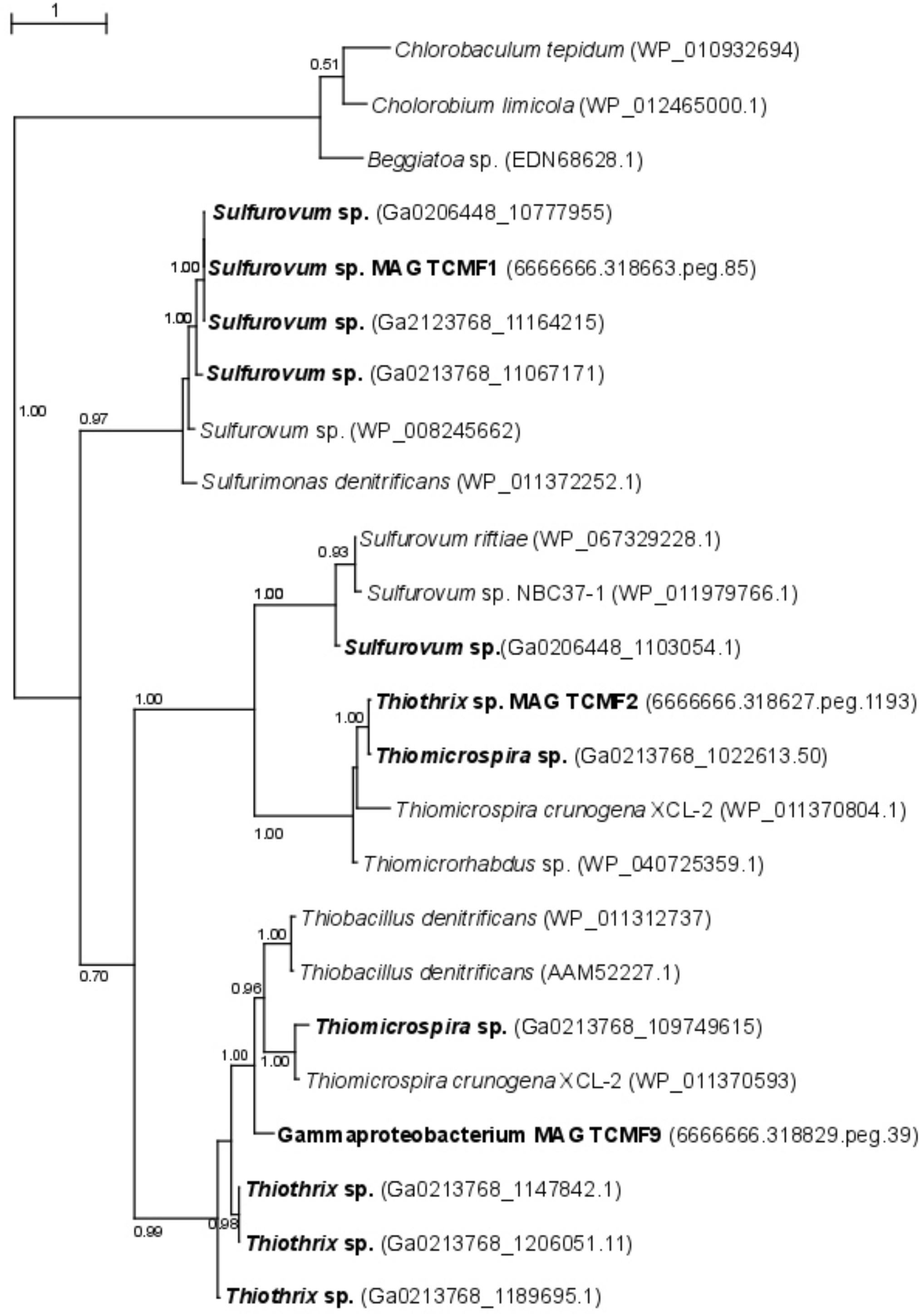
Maximum likelihood phylogenetic tree showing the position of the sulfide quinone oxireductase amino acid sequences, SQR, obtained in this study (indicated in boldface). Closely related sequences were obtained from GenBank. Approximate likelihood ratio test (aLRT) values for branch support are indicated. Bar, 1% estimated substitutions.

## Discussion

Chemosynthetic microbial biofilm are commonly found in geothermal and sulfidic environments (Sievert and Vetriani, 2012), including cold seeps (Ristova *et al*., 2015), sulfidic caves (Macalady *et al*., 2006), thermal springs (Beam *et al*., 2016), mud volcanoes (Heijs *et al*., 2005) as well as deep-sea (Gulmann et al., 2015, O’Brien et al., 2015) and shallow-water hydrothermal vents (Miranda *et al*., 2016; Patwardhan *et al*., 2018). Members of *Gammaproteobacteria* and *Epsilonproteobacteria* dominate most of these biofilms and a spatial segregation between the two classes of bacteria has been correlated with the *in situ* sulfide concentration (Engel *et al*., 2004; Crépeau *et al*., 2011; Gulmann *et al*., 2015; O’Brien *et al*., 2015; Miranda *et al*., 2016; Macalady et al., 2008; Meier et al., 2017).

At Tor Caldara, a submarine gas vent in the Tyrrhenian Sea characterized by vigorous venting of CO_2_ and unusually a high concentration of H_2_S (avg. of 23.13 mol%; Patwardhan *et al*., 2018), white filamentous microbial communities grow profusely on the rocks near the gas venting. To investigate the composition and function of established and young biofilms, we collected native substrates (EF) as well as glass slides (YF) deployed in the vicinity of a vent. Within 4 days, filamentous biofilms colonized the slides.

Our survey of 16S rRNA genes and transcripts revealed a shift in the two main taxonomic groups during colonization, whereby sulfur-oxidizing *Epsilonproteobacteria* dominated the YF biofilms, while *Gammaproteobacteria* became prevalent in the EF community. Further, we showed that representative species of *Epsilon*- and *Gammaproteobacteria* are adapted to different sulfide concentrations (Patwardhan *et al*., 2018). In this study, we integrated metagenomic and metaproteomic approaches to investigate the *in situ* expression of central metabolic pathways related to carbon fixation and energy conservation in the YF and EF biofilms. In line with our previous 16S rRNA amplicon survey, the metagenome-extracted 16S rRNA genes showed that *Epsilonproteobacteria* and *Gammaproteobacteria* dominated the YF and EF biofilms, respectively (Figs. 1a and b).

### The relative expression of the rTCA and CBB cycles for carbon fixation reflects the taxonomic composition of the YF and EF biofilms

Chemoautotrophs are the primary producers at hydrothermal vents, forming the basis of the food web by fixing inorganic CO_2_ into organic material (Jannasch and Wirsen, 1979; Karl *et al*., 1980, Sievert and Vetriani, 2012). Till date, six pathways for carbon fixation are known: the Calvin-Benson-Bassham (CBB) reductive pentose phosphate cycle, the reductive tricarboxylic acid cycle (rTCA), the reductive acetyl coenzyme A pathway(Wood-Ljungdahl), the 3-hydroxypropionate bi-cycle, the 3-hydroxypropionate/4-hydroxybutyrate cycle, and the dicarboxylate/4-hydroxybutyrate cycle (Nakagawa and Takai, 2008; Berg, 2011). The rTCA and CBB cycles are most prevalent at diffuse-flow hydrothermal vents and, in these habitats, they are generally diagnostic of *Epsilon*- and *Gammaproteobacteria*, respectively (Nakagawa and Takai, 2008). Enrichment of proteins of the rTCA cycle affiliated with *Epsilonproteobacteria*, in addition to a similar signature at the gene level, confirmed that autotrophy using the rTCA cycle in *Epsilonproteobacteria* was the predominant mechanism for carbon fixation in the YF communities (Figs. 2, 6 and Suppl. Tables 1, 2). The majority of the rTCA genes and proteins were classified as *Sulfurovum* spp. (Fig. 2), which is known to be ubiquitous in vent environments (Inagaki, 2004; Dahle *et al*., 2013; Mino *et al*., 2014; Giovannelli *et al*., 2016). These findings underline the importance of *Sulfurovum* spp. in the YF community and are in accordance with data from the 16S rRNA diversity analysis (Patwardhan *et al*., 2018; Fig. 1). While the EF community also encoded genes and expressed enzymes of the rTCA cycle, the CBB cycle was overrepresented in the EF biofilm at both the gene and protein levels, with contributions from various genera within the *Gammaproteobacteria*, in particular *Thiomicrospira, Thioalkovibrio* and *Thiothrix-related* spp. (Figs. 2, 6 and Suppl. Tables 1, 2). This is also in line with our 16S rRNA survey, which showed these genera as dominant in the EF biofilm (Patwardhan *et al*., 2018; Fig. 1). Thus, we observed a shift from rTCA-based carbon fixation (*Epsilonproteobacteria*) to CBB-based carbon fixation (*Gammaproteobacteria*) during biofilm maturation. The lower energy demand of the rTCA cycle, compared to the CBB cycle (Nakagawa and Takai, 2008; Hügler and Sievert, 2011), might provide an advantage to the *Epsilonproteobacteria* during early substrate colonization.

**Fig. 6.**
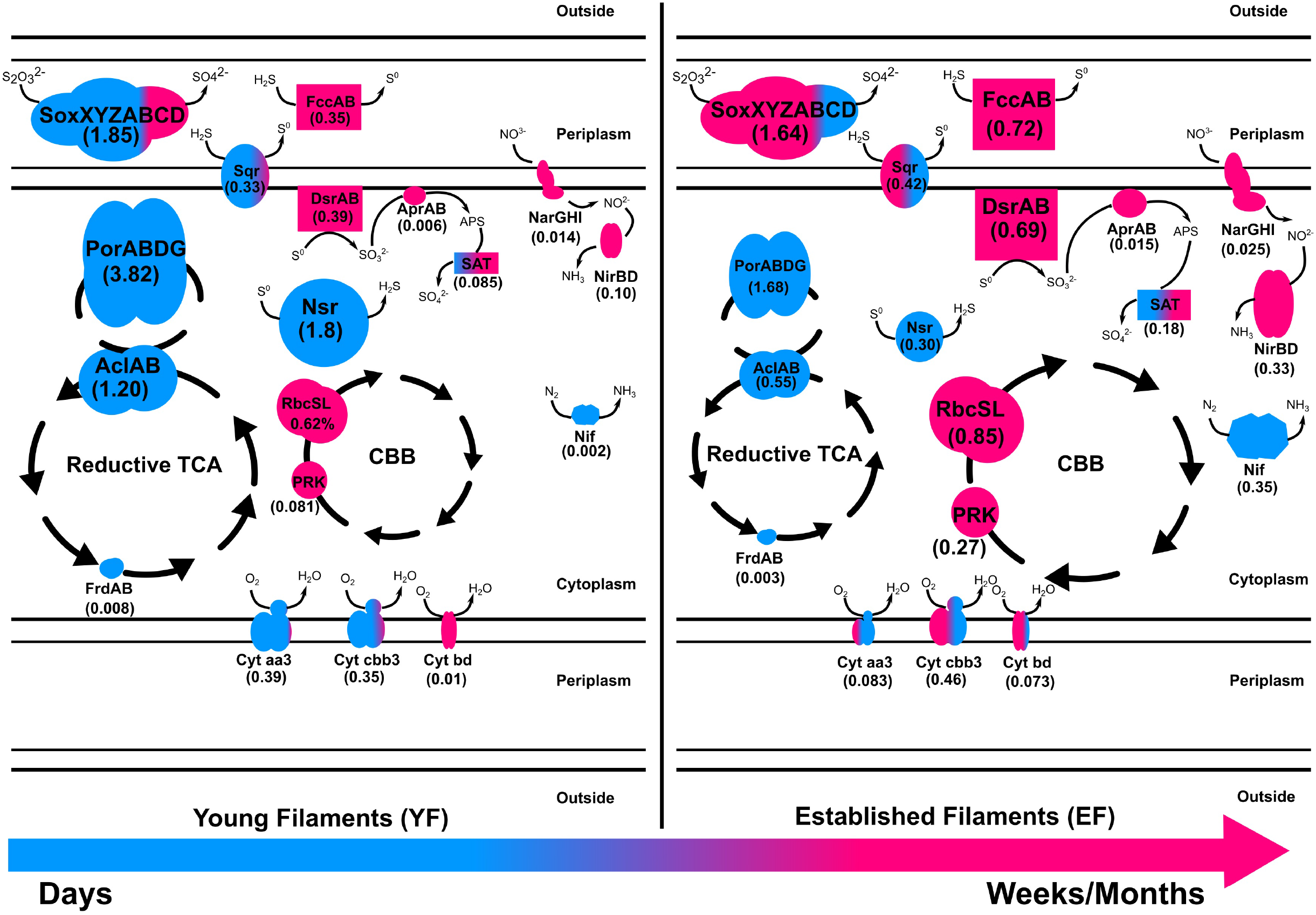
Metaproteomic-based reconstruction of the central metabolic pathways expressed by the Tor Caldara young and established biofilm communities. The size of the enzymatic complexes is proportional to their abundance in the metaproteome. The numbers indicate the normalized protein abundance (%). The colors indicate the taxonomic affiliation of the enzymes and reflect the relative proportions of the two predominant bacterial classes. Blue: *Epsilonproteobacteria;* Pink: *Gammaproteobacteria*.

### The expression of different pathways for sulfide/sulfur oxidation in the YF and EF biofilms may reflect adaptations to fluctuating redox regimes

In shallow-water geothermal systems, the concentration of reduced sulfur compounds and other electron donors for microbial oxidations, along with the availability of oxidants, vary considerably over time (Yücel *et al*, 2013; Price and Giovannelli, 2017). Hence, chemolithoautotrophic microorganisms have adapted to such fluctuating redox regimes. In particular, reduced sulfur compounds represent an important energy source that drives energy-yielding reactions in geothermal environments (Nakagawa and Takai, 2008). The abundance and range of various genes and proteins involved in sulfur oxidation pathways presented here validated our initial findings about the importance of sulfur-oxidizing bacteria at Tor Caldara (Patwardhan *et al*., 2018). For instance, the Sox pathway for sulfur oxidation was one of the most abundant energy-yielding pathway recovered both in the metagenomes and metaproteomes (Figs. 3, 4, 6 and Suppl. Tables 1, 2).

Thiosulfate oxidation catalyzed by the multienzyme complex Sox is widespread in sulfur-oxidizing chemoautotrophs (Friedrich *et al*., 2001, 2005; Yamamoto and Takai, 2011). This enzyme complex, consisting of four protein components, soxYZ, soxXA, soxB and soxCD, catalyzes the oxidation of thiosulfate and sulfide to sulfate. The abundance of the Sox complex in both filament communities at the gene as well as the protein (YF:1.8%; EF:1.6%) level indicates that this is the predominant pathway used by the biofilm bacteria to conserve energy (Figs. 3a, 4a and Suppl. Tables 1, 2). This is consistent with a metaproteogenomic study of chimney-associated microbial communities from deep-sea hydrothermal vents, which revealed that Sox represented one of the most highly expressed energy-yielding pathways (Pjevac *et al*., 2018). However, the taxonomic affiliation of the *sox* reads (Fig. 3b) and Sox enzymes (Fig. 4b) from the Tor Caldara biofilms shows a finer resolution, indicating that *Epsilonproteobacteria* are mostly expressing the Sox complex in the YF biofilm, while *Gammaproteobacteria* take over that same role in the EF biofilms (Fig. 6).

A similar trend is evident in the profile of the sulfide:quinone oxidoreductase enzyme complex (SQR) involved in oxidation of sulfide to elemental sulfur, which is predominantly expressed by *Epsilonproteobacteria* in the YF filaments, while its contribution by *Gammaproteobacteria* increases in the EF filament community (Fig. 4b and 6). In addition to SQR, in some chemolithotrophic bacteria oxidation of hydrogen sulfide to elemental sulfur is catalyzed by the flavocytochrome *c*-sulfide dehydrogenase (FccAB; Visser *et al*., 1997; Mußmann *et al*., 2007). The metaproteomic data presented here show that the FccAB enzyme is affiliated only with *Gammaproteobacteria*, and that its expression is higher in the EF biofilm than in the YF community (Figs. 4a and 6). The bioenergetics of microbial sulfide oxidation might provide a framework to interpret the SQR and Fcc expression profiles in the Tor Caldara biofilms. Since the midpoint potential of the NAD^+^/NADH couple at pH 7 is approximately 50 mV more negative than the midpoint potential of the S^0^/H_2_S couple, chemolithotrophic SOBs require energy to transport electrons from hydrogen sulfide upwards to NAD^+^ by reverse electron flow. In SQR-catalyzed sulfide oxidation, the electrons from sulfide enter the transport chain at the level of quinones, generating the necessary electrochemical proton potential across the membrane for reverse electron transfer (Griesbeck *et al*. 2000). However, when Fcc is involved, the electrons enter the transport chain at the level of *c*-type cytochromes, whose midpoint potential is more electropositive than that of quinones. That increases the energetic burden for the reverse transfer of electrons upwards to NAD^+^ and implies that, in chemolithotrophic SOBs, sulfide oxidation by SQR provides more energy than sulfide oxidation by Fcc (Griesbeck *et al*. 2000). However, due to its high affinity for sulfide (Brune, 1995), Fcc is hypothesized to be expressed at low sulfide concentrations, supplementing the energetically more efficient SQR. Previous work on the Tor Caldara biofilms demonstrated that *Epsilonproteobacteria* are adapted to higher sulfide concentrations than *Gammaproteobacteria* and that they are the pioneer colonizers at this site (Patwardhan *et al*., 2018). We hypothesize that, early in the colonization process (YF biofilm), *Epsilonproteobacteria* might be driving down sulfide levels within the biofilms and that the increased expression of the high affinity FccAB in EF might provide an advantage to the *Gammaproteobacteria* in the subsequent stages of colonization (EF biofilm; Fig. 6). The hypothesis that Fcc is more prevalent under low sulfide and more oxidized conditions (Griesbeck *et al*.; Brune, 1995) supports our model. Overall, the co-expression of the energy efficient SQR and the high affinity Fcc may provide metabolic flexibility in response to fluctuating sulfide concentrations within the biofilm communities.

The reverse dissimilatory sulfate reductase pathway (rDSR) oxidizes stored intracellular sulfur to sulfate via sulfite (Friedrich *et al*., 2005). Consistent with findings in other filamentous *Gammaproteobacteria* (Mußmann *et al*., 2007; Chernousova *et al*., 2009; Sharrar *et al*., 2017), all three proteins involved in this pathway, namely, DsrAB, AprAB and SAT, were affiliated with *Gammaproteobacteria* and were enriched in the EF biofilm (Figs. 4, 6 and Suppl. Table 2). The expression of the rDSR pathway is in accordance with the stored intracellular elemental sulfur observed in electron micrographs of EF (Patwardhan *et al*., 2018), while its higher expression in the EF biofilm suggests that this community may experience transient depletions of hydrogen sulfide. This stored sulfur might also be disproportionated into sulfite, thiosulfate and sulfide as evidenced by the expression of gammaproteobacterial Sor (Fig. 4) in the EF biofilm, and might provide a competitive advantage to the secondary colonizers (Kletzin, 1989; Veith *et al*., 2012). Since the Dsr pathway for S-oxidation is not encoded in the currently available genomes of *Epsilonproteobacteria* (Yamamoto and Takai, 2011), the epsiloproteobacterial SAT found in the metaproteome is probably involved in sulfate assimilation.

Sulfur oxidation was complemented with sulfur reduction evidenced by high expression of the a putative *Sulfurovum-affiliated* NADH-dependent sulfur reductase (NSR; Schut *et al*., 2007) in the YF (Figs. 4, 6 and Suppl. Table 2). Despite sulfur respiration is known to be coupled to hydrogen oxidation in two *Sulfurovum* spp. (Yamamoto Masahiro *et al*., 2010; Mino *et al*., 2014), we did not detect hydrogenases in the metaproteomes. Since hydrogen concentrations are low at Tor Caldara, the *Sulfurovum* spp. found there may be adapted to oxidize alternative energy sources, such as malate and/or formate (Campbell *et al*., 2006; Yamamoto Masahiro *et al*., 2010; Keller *et al*., 2015).

### *Epsiloproteobacteria-mediated* nitrogen fixation and *Gammaproteobacteria-mediated* nitrate reduction are prevalent in the EF biofilm

Microbial nitrate respiration occurs via two main pathways, dissimilatory nitrate reduction to ammonia (DNRA) and denitrification. Both processes have been investigated in marine geothermal environments (Bourbonnais *et al*., 2012; Bourbonnais *et al*., 2012; Pérez-Rodríguez *et al*., 2013; Vetriani *et al*., 2014). The first step in nitrate reduction is carried out by the enzyme nitrate reductase. Currently, two different types of respiratory nitrate reductases, the membrane-bound Nar and the periplasmic Nap, are known to occur in bacteria (Richardson *et al*., 2001). Nar is a low-affinity enzyme expressed under nitrate rich conditions, whereas Nap is a high-affinity enzyme whose expression is favored under low nitrate conditions (Potter *et al*., 1999; Mintmier *et al*., 2018). The genes encoding both Nar and Nap were present in the EF and YF communities, as well as the genes encoding various nitrite reductases (NirBD, NirK, NirS and NrfA), the nitric oxide reductase (NorB) and the nitrous oxide reductase (NosZ; (Fig. 3a and Suppl. Table 1). However, metaproteomic analyses showed that only the *Gammaproteobacteria*-affiliated Nar and Nir enzymes were expressed, while Nap and the enzymes involved in the denitrification pathway could not detected (Figs. 4b). The coastal area at Tor Caldara is likely enriched in nitrate (possibly from runoff of lawn fertilizers), which might explain the expression of NarGHI, but not of NapAB (Fig 4a and Suppl. Table 2). The NirBD nitrite reductase, often involved in the reduction of nitrite to ammonia in DNRA (Reyes *et al*., 2017), was also found to be abundant. Both the Nar and Nir enzymes were enriched in the EF biofilm, indicating that nitrate respiration is prevalent in the established community.

Apart from nitrate respiration, nitrogen fixation also plays an important role in nitrogen cycling at vents (Rau, 1981; Mehta *et al*., 2003). Expression of epsilonproteobacterial NifDHK was observed in both filament communities but was highly enriched in the EF (Figs. 4, 6 and Suppl. Table 2). While nitrogen fixation is not common in *Epsilonproteobacteria*, recent studies have shown that some members of this class have the genomic potential for it (Keller *et al*., 2015; Waite *et al*., 2017), and the ability to fix nitrogen was demonstrated experimentally in *Lebetimonas* spp. (Meyer and Huber, 2013). Nitrogen fixation is an energy expensive reaction and nitrogenases are usually expressed in nitrogen limiting conditions (Olivares *et al*., 2013). We hypothesize that, when the filamentous community transitions from a young to an established state, nitrate respiration carried out by *Gammaproteobacteria* might result in a depletion in nitrogen within the biofilm microenvironment. Hence, the ability to fix nitrogen might give the *Epsilonproteobacteria* an advantage while competing for a fixed nitrogen source against the secondary colonizers in the established biofilm.

### Enzymes for aerobic and microaerobic respiration are differentially expressed in the YF and EF biofilms

Most of the well-characterized bacterial oxidases belong to the heme-copper superoxidase family. There are two types of oxidases that use cytochrome *c* as a substrate: cytochrome *c* oxidase aa3-type and cytochrome *c* oxidase cbb3-type. The former is expressed in aerobic (high oxygen) conditions while the latter is expressed in microaerobic (low oxygen) conditions. Additionally, cytochrome bd, a ubiquinol oxidase, is also expressed in microaerobic conditions (García-Horsman *et al*., 1994; Pitcher and Watmough, 2004). At Tor Caldara, the microaerobic, high affinity cbb3-type and bd-type cytochrome oxidase were enriched in EF biofilm and were mostly classified as *Gammaproteobacteria* (Figs. 4, 6 and Suppl. Table 2), suggesting that the biofilms experience microaeobic/anoxic conditions as they mature. On the other hand, primary colonizers in the YF might experience oxic to micro-aerobic conditions as suggested by equal abundance of aerobic aa3 type and micro-aerobic cbb3 type cytochrome *c* oxidase classified dominantly as *Epsilonproteobacteria*.

### Overall view: A metaproteome-based model of microbial colonization at the Tor Caldara gas vents

Based on the metaproteomic data, we reconstructed the central metabolic pathways expressed by the Tor Caldara young and established biofilm communities (Fig. 6). According to this model, substrates exposed to the gas emissions are initially colonized by a population dominated by sulfide-tolerant *Epsilonproteobacteria* (Fig. 1; Patwardhan et al., 2018). Members of this class oxidize various reduced sulfur compounds via the Sox and SQR pathways coupled to oxygen reduction (both aerobic and micro-aerobic), while fixation of carbon dioxide occurs via the rTCA cycle. In the early stages of colonization, *Gammaproteobacteria* are present, but underrepresented (Figs. 1 and 6). Over time, sulfide oxidation by *Epsilonproteobacteria* may lower the sulfide concentration within the biofilm community, conditioning the environment for the growth of the less sulfide-tolerant *Gammaproteobacteria* (Patwardhan *et al*., 2018), which dominate the established filaments. *Gammaproteobacteria* also oxidize various sulfur compounds using the Sox and SQR pathways, but this reaction is now coupled to either microaerobic and/or anaerobic respiration using oxygen and nitrate as terminal electron acceptors, respectively. The high affinity, periplasmic FccAB enzyme is co-expressed with SQR in response to fluctuating concentrations of sulfide, while elemental sulfur is stored in the cytoplasm. Energy thus conserved is used to fix carbon dioxide via the CBB cycle (Fig. 6). When sulfide availability becomes limiting within the biofilm community, certain *Gammaproteobacteria* further oxidize the stored sulfur to sulfate via the reverse dissimilatory sulfate reductase pathway, thus making full use of their metabolic repertoire. As the filamentous community transitions from a young to a more established stage, anaerobic respiration by the *Gammaproteobacteria* may lead to loss of nitrate from the system. This triggers increased nitrogen fixation by the *Epsilonpreoteobacteria* as a potential strategy to compete with the *Gammaproteobacteria*.

## Conclusion

Our study revealed a complex pattern of protein expression in chemosynthetic biofilm communities that colonize the gas vent system at Tor Caldara. Functions common to chemosynthetic microbial communities, such as carbon fixation and sulfide oxidation, are catalyzed via the expression of different enzymatic complexes encoded by two predominant groups of SOBs belonging to the *Epsilon*- and *Gammaproteobacteria*. We hypothesize that the expression of these enzymatic complexes is finely tuned to the chemical and physical conditions within the biofilm communities, and reflect the physiological characteristics of the two groups of SOBs. Overall, the enzyme expression profiles obtained in this work reveal metabolic flexibility and adaptations to fluctuating redox regimes during colonization of the shallow-water gas vents of Tor Caldara. These results are consistent with the previously reported transition from *Epsilonproteobacteria* - the pioneer colonists - to *Gammaproteobacteria* at Tor Caldara.

## Acknowledgements

We thank the crew of the R/V *Antares* for their support during the sampling operations, and Satyajit Rajapurkar for help and guidance with the bioinformatics analyses. This work was partially supported by the following grants to CV: NSF OCE 11-24141, NSF OCE 11-36451, NSF MCB 15-17567 and NASA NNX15AM18G. This paper is C-DEBI contribution xx.

The authors declare that they have no conflict of interest.

---

**Supplementary Table 1.** Metagenomes: Normalized abundances of key genes involved in carbon fixation, nitrogen & sulfur metabolism, oxygen respiration, and heavy metal detoxification pathways expressed as transcripts (reads) per million (TPM).

| Gene | TPM |  |
| --- | --- | --- |
|  | Young Filaments (YF) | Established Filaments (EF) |
| ATP-citrate lyase (aclAB) | 681.5 | 210.34 |
| Fumarate reductase (frdAB) | 1331.01 | 115.24 |
| Pyruvate ferredoxin oxidoreductase (porABDG) | 1141.07 | 550.12 |
| Phosphoribulokinase (prk) | 99.47 | 207.22 |
| Ribulose-bisphosphate carboxylase (rbcSL) | 182.77 | 408.48 |
| Nitrate reductase (napAB) | 381.01 | 153.62 |
| Nitrate reductase (narGHI) | 21.55 | 86.98 |
| Nitric oxide reductase (norB) | 1.35 | 3.72 |
| Nitrite reductase (nirBD) | 312.33 | 498.28 |
| Nitrite reductase, cytochrome c-552 (nrfA) | 0.37 | 4.92 |
| Nitrite reductase, NO-forming (nirK) | 0.77 | 7.69 |
| Nitrite reductase, NO-forming (nirS) | 2.23 | 4.81 |
| Nitrogenase (nifDHK) | 499.47 | 325.34 |
| Nitrous-oxide reductase (nosZ) | 6.71 | 12.34 |
| Adenylylsulfate reductase (aprAB) | 17.5 | 26.49 |
| Polysulfide reductase (psrA) | 2.61 | 11.83 |
| Sox system (soxXYZABCD) | 2553.8 | 2308.16 |
| Sulfate adenylyltransferase (sat) | 237.44 | 255.85 |
| Sulfide dehydrogenase (fccAB) | 635.6 | 1402.89 |
| Sulfide:quinone oxidoreductase (sqr) | 375.84 | 843.84 |
| Sulfide dehydrogenase | 52.65 | 2.22 |
| Sulfite reductase (dsrAB) | 76.72 | 210.2 |
| Sulfur oxygenase/reductase (sor) | 0 | 38.43 |
| Putative Sulfur reductase (nsr) | 323.84 | 394.22 |
| Cytochrome c oxidase, aa3-type (coxABCD) | 834.02 | 665.43 |
| Cytochrome c oxidase, cbb3-type (ccoNOPQ) | 944.63 | 1026.12 |
| Cytochrome bd ubiquinol oxidase (cydAB) | 284.87 | 392.7 |
| Arsenate reductase (arsC) | 527.49 | 543.82 |
| Mercuric reductase (merA) | 502.49 | 174.16 |
| Selenate detoxification (dedA) | 0.69 | 15.14 |

**Supplementary Table 2.** Metaproteomes: Normalized abundances of key proteins involved in carbon fixation, nitrogen & sulfur metabolism, oxygen respiration, and heavy metal detoxification pathways expressed as a percentage of total proteins.

| Protein | Percent |  |
| --- | --- | --- |
|  | Young Filaments (YF) | Established Filaments (EF) |
| ATP-citrate lyase (AclAB) | 1.202 | 0.545 |
| Fumarate reductase (FrdAB) | 0.008 | 0.003 |
| Pyruvate ferredoxin oxidoreductase (PorABDG) | 3.825 | 1.677 |
| Phosphoribulokinase (PRK) | 0.081 | 0.267 |
| Ribulose-bisphosphate carboxylase (RbcSL) | 0.623 | 0.846 |
| Nitrate reductase (NarGHI) | 0.014 | 0.025 |
| Nitrite reductase (NirBD) | 0.103 | 0.329 |
| Nitrogenase (NifDHK) | 0.002 | 0.354 |
| Adenylylsulfate reductase (AprAB) | 0.006 | 0.015 |
| Sox system (SoxXYZABCD) | 1.85 | 1.635 |
| Sulfate adenylyltransferase (SAT) | 0.085 | 0.18 |
| Sulfide dehydrogenase (FccAB) | 0.351 | 0.72 |
| Sulfide:quinone oxidoreductase (Sqr) | 0.336 | 0.42 |
| Sulfite reductase (DsrAB) | 0.399 | 0.639 |
| Sulfur oxygenase/reductase (Sor) | 0.013 | 0.06 |
| Putative Sulfur reductase (Nsr) | 1.799 | 0.301 |
| Cytochrome c oxidase, aa3-type (CoxABCD) | 0.397 | 0.083 |
| Cytochrome c oxidase, cbb3-type (CcoNOPQ) | 0.357 | 0.469 |
| Cytochrome bd ubiquinol oxidase (CydAB) | 0.014 | 0.073 |
| Arsenate reductase (ArsC) | 0 | 0.003 |
| Selenate detoxification (DedA) | 0.008 | 0.004 |

## References

Anders, S., Pyl, P.T., and Huber, W. (2015) HTSeq—a Python framework to work with high-throughput sequencing data. Bioinformatics 31: 166–169.

Anisimova, M. and Gascuel, O. (2006) Approximate Likelihood-Ratio Test for Branches: A Fast, Accurate, and Powerful Alternative. Syst Biol 55: 539–552.

Barry, P.H., de Moor, J.M., Giovannelli, D., Schrenk, M., Hummer, D., Lopez, T., Pratt, C.A., Alpízar Segura, Y., Battaglia, A., P. Beaudry, P., Bini, G., Cascante, M., d’Errico, G., di Carlo, M., Fattorini, D., Fullerton, K., Gazel, E., González, G., Halldórsson, S. A., Iacovino, K., Kulongoski, J.T., Manini, E., Martínez, M., Miller, H., Nakagawa, M., Ono, S., Patwardhan, S., Ramírez, C.J., Regoli, F., Smedile, F., Turner, S., Vetriani, C., Yücel, M., Ballentine, C.J., Fischer, T.P., Hilton, D.R., Lloyd. K.G. (2019). Forearc carbon sequestration reduces long-term volatile recycling into the mantle. Nature 568: 487–492.

Beam, J.P., Bernstein, H.C., Jay, Z.J., Kozubal, M.A., Jennings, R. deM., Tringe, S.G., and Inskeep, W.P. (2016) Assembly and Succession of Iron Oxide Microbial Mat Communities in Acidic Geothermal Springs. Front Microbiol 7: 25. DOI=10.3389/fmicb.2016.00025

Bengtsson-Palme Johan, Hartmann Martin, Eriksson Karl Martin, Pal Chandan, Thorell Kaisa, Larsson Dan Göran Joakim, and Nilsson Rolf Henrik (2015) metaxa2: improved identification and taxonomic classification of small and large subunit rRNA in metagenomic data. Molecular Ecology Resources 15: 1403–1414.

Berg, I.A. (2011) Ecological Aspects of the Distribution of Different Autotrophic CO2 Fixation Pathways. Appl Environ Microbiol 77: 1925–1936.

Boden, R., Scott, K.M., Williams, J., Russel, S., Antonen, K., Rae, A.W., and Hutt, L.P. (2017) An evaluation of Thiomicrospira, Hydrogenovibrio and Thioalkalimicrobium: reclassification of four species of Thiomicrospira to each Thiomicrorhabdus gen. nov. and Hydrogenovibrio, and reclassification of all four species of Thioalkalimicrobium to Thiomicrospira. International Journal of Systematic and Evolutionary Microbiology 67: 1140–1151.

Bourbonnais, A., Juniper, S.K., Butterfield, D.A., Devol, A.H., Kuypers, M.M.M., Lavik, G., et al. (2012) Activity and abundance of denitrifying bacteria in the subsurface biosphere of diffuse hydrothermal vents of the Juan de Fuca Ridge. Biogeosciences 9: 4661–4678.

Bourbonnais, A., Lehmann, M.F., Butterfield, D.A., and Juniper, S.K. (2012) Subseafloor nitrogen transformations in diffuse hydrothermal vent fluids of the Juan de Fuca Ridge evidenced by the isotopic composition of nitrate and ammonium: nitrogen cycle in hydrothermal vents. Geochemistry, Geophysics, Geosystems 13: Q02T01, doi:10.1029/2011GC003863.

Bowers, R., Kyrpides, N., Stepanauskas, R. et al. (2017) Minimum information about a single amplified genome (MISAG) and a metagenome-assembled genome (MIMAG) of bacteria and archaea. Nat Biotechnol 35: 725–731.

Brune, D.C. (1995) Sulfur Compounds as Photosynthetic Electron Donors. In, Anoxygenic Photosynthetic Bacteria, Advances in Photosynthesis and Respiration. Springer, Dordrecht, pp. 847–870.

Campbell, B.J., Engel, A.S., Porter, M.L., and Takai, K. (2006) The versatile ϵ-proteobacteria: key players in sulphidic habitats. Nat Rev Micro 4: 458–468.

Carapezza, M.L., Barberi, F., Ranaldi, M., Ricci, T., Tarchini, L., Barrancos, J., et al. (2012) Hazardous gas emissions from the flanks of the quiescent Colli Albani volcano (Rome, Italy). Applied Geochemistry 27: 1767–1782.

Carapezza, M.L. and Tarchini, L. (2007) Accidental gas emission from shallow pressurized aquifers at Alban Hills volcano (Rome, Italy): Geochemical evidence of magmatic degassing? Journal of Volcanology and Geothermal Research 165: 5–16.

Chernousova, E., Gridneva, E., Grabovich, M., Dubinina, G., Akimov, V., Rossetti, S., and Kuever, J. (2009) Thiothrix caldifontis sp. nov. and Thiothrix lacustris sp. nov., gammaproteobacteria isolated from sulfide springs. International Journal of Systematic and Evolutionary Microbiology 59: 3128–3135.

Craig, R. and Beavis, R.C. (2004) TANDEM: matching proteins with tandem mass spectra. Bioinformatics 20: 1466–1467.

Crépeau, V., Cambon Bonavita, M.-A., Lesongeur, F., Randrianalivelo, H., Sarradin, P.-M., Sarrazin, J., and Godfroy, A. (2011) Diversity and function in microbial mats from the Lucky Strike hydrothermal vent field. FEMS Microbiol Ecol 76: 524–540.

Dahle, H., Roalkvam, I., Thorseth, I.H., Pedersen, R.B., and Steen, I.H. (2013) The versatile in situ gene expression of an Epsilonproteobacteria-dominated biofilm from a hydrothermal chimney. Environmental Microbiology Reports 5: 282–290.

Dando, P.R., Stüben, D., and Varnavas, S.P. (1999) Hydrothermalism in the Mediterranean Sea. Progress in Oceanography 44: 333–367.

Engel, A.S., Porter, M.L., Stern, L.A., Quinlan, S., and Bennett, P.C. (2004) Bacterial diversity and ecosystem function of filamentous microbial mats from aphotic (cave) sulfidic springs dominated by chemolithoautotrophic “Epsilonproteobacteria.” FEMS Microbiol Ecol 51: 31–53.

Fisher, T.P. (2008). Fluxes of volatiles (H_2_O, CO_2_, N_2_, Cl, F) from arc volcanoes. Geochem J 42: 21–38.

Friedrich, C.G., Bardischewsky, F., Rother, D., Quentmeier, A., and Fischer, J. (2005) Prokaryotic sulfur oxidation. Current Opinion in Microbiology 8: 253–259.

Friedrich, C.G., Rother, D., Bardischewsky, F., Quentmeier, A., and Fischer, J. (2001) Oxidation of Reduced Inorganic Sulfur Compounds by Bacteria: Emergence of a Common Mechanism? Appl Environ Microbiol 67: 2873–2882.

García-Horsman, J.A., Barquera, B., Rumbley, J., Ma, J., and Gennis, R.B. (1994) The superfamily of heme-copper respiratory oxidases. J Bacteriol 176: 5587–5600.

Giovannelli, D., Chung, M., Staley, J., Starovoytov, V., Le Bris, N., and Vetriani, C. (2016) Sulfurovum riftiae sp. nov., a mesophilic, thiosulfate-oxidizing, nitrate-reducing chemolithoautotrophic epsilonproteobacterium isolated from the tube of the deep-sea hydrothermal vent polychaete Riftia pachyptila. Int J Syst Evol Microbiol 66: 2697–2701.

Giovannelli, D., d’Errico, G., Manini, E., Yakimov, M.M., and Vetriani, C. (2013) Diversity and phylogenetic analyses of bacteria from a shallow-water hydrothermal vent in Milos island (Greece). Front Microbiol 4: 184.

Griesbeck, C., Hauska, G., and Schütz, M. (2000) Biological Sulfide Oxidation: Sulfide-Quinone Reductase (SQR), the Primary Reaction. In: Pandalai, S.G. (ed): Recent Research Developments in Microbiology, Vol. 4, 179–203. Research Signpost, Trivadrum, India.

Gugliandolo, C. and Maugeri, T.L. (1993) Chemolithotrophic, sulfur-oxidizing bacteria from a marine, shallow hydrothermal vent of Vulcano (Italy). Geomicrobiology Journal 11: 109–120.

Guindon, S., Dufayard, J.-F., Lefort, V., Anisimova, M., Hordijk, W., and Gascuel, O. (2010) New Algorithms and Methods to Estimate Maximum-Likelihood Phylogenies: Assessing the Performance of PhyML 3.0. Syst Biol 59: 307–321.

Gulmann, L.K., Beaulieu, S.E., Shank, T.M., Ding, K., Seyfried, W.E., and Sievert, S.M. (2015) Bacterial diversity and successional patterns during biofilm formation on freshly exposed basalt surfaces at diffuse-flow deep-sea vents. Front Microbiol 6: 901.

Gurevich, A., Saveliev, V., Vyahhi, N., and Tesler, G. (2013) QUAST: quality assessment tool for genome assemblies. Bioinformatics 29: 1072–1075.

Heijs, S.K., Sinninghe DamstÃ©, J.S., and Forney, L.J. (2005) Characterization of a deep-sea microbial mat from an active cold seep at the Milano mud volcano in the Eastern Mediterranean Sea. FEMS Microbiology Ecology 54: 47–56.

Hügler, M. and Sievert, S.M. (2011) Beyond the Calvin Cycle: Autotrophic Carbon Fixation in the Ocean. Annual Review of Marine Science 3: 261–289.

Inagaki, F. (2004) Sulfurovum lithotrophicum gen. nov., sp. nov., a novel sulfur-oxidizing chemolithoautotroph within the -Proteobacteria isolated from Okinawa Trough hydrothermal sediments. International Journal Of Systematic And Evolutionary Microbiology 54: 1477–1482.

Jannasch, H.W. and Wirsen, C.O. (1979) Chemosynthetic Primary Production at East Pacific Sea Floor Spreading Centers. BioScience 29: 592–598.

Kalanetra, K.M., Huston, S.L., and Nelson, D.C. (2004) Novel, Attached, Sulfur-Oxidizing Bacteria at Shallow Hydrothermal Vents Possess Vacuoles Not Involved in Respiratory Nitrate Accumulation. Appl Environ Microbiol 70: 7487–7496.

Karl, D.M. (Univ of H., Wirsen, C.O., and Jannasch, H.W. (1980) Deep-sea primary production at the Galapagos hydrothermal vents. Science; (United States) 207:.

Keller, A.H., Schleinitz, K.M., Starke, R., Bertilsson, S., Vogt, C., and Kleinsteuber, S. (2015) Metagenome-Based Metabolic Reconstruction Reveals the Ecophysiological Function of Epsilonproteobacteria in a Hydrocarbon-Contaminated Sulfidic Aquifer. Front Microbiol 6:.

Kerfahi, D., Hall-Spencer, J.M., Tripathi, B.M., Milazzo, M., Lee, J., and Adams, J.M. Shallow Water Marine Sediment Bacterial Community Shifts Along a Natural CO2 Gradient in the Mediterranean Sea Off Vulcano, Italy. Microb Ecol 1–10.

Kletzin, A. (1989) Coupled enzymatic production of sulfite, thiosulfate, and hydrogen sulfide from sulfur: purification and properties of a sulfur oxygenase reductase from the facultatively anaerobic archaebacterium Desulfurolobus ambivalens. J Bacteriol 171: 1638–1643.

Langmead, B., Trapnell, C., Pop, M., and Salzberg, S.L. (2009) Ultrafast and memory-efficient alignment of short DNA sequences to the human genome. Genome Biology 10: R25.

Le, S.Q. and Gascuel, O. (2008) An Improved General Amino Acid Replacement Matrix. Mol Biol Evol 25: 1307–1320.

Ledgham, F., Quest, B., Vallaeys, T., Mergeay, M., and Covès, J. (2005) A probable link between the DedA protein and resistance to selenite. Res Microbiol 156: 367–374.

Li, D., Liu, C.-M., Luo, R., Sadakane, K., and Lam, T.-W. (2015) MEGAHIT: an ultra-fast single-node solution for large and complex metagenomics assembly via succinct de Bruijn graph. Bioinformatics 31: 1674–1676.

Macalady, J.L., Lyon, E.H., Koffman, B., Albertson, L.K., Meyer, K., Galdenzi, S., and Mariani, S. (2006) Dominant Microbial Populations in Limestone-Corroding Stream Biofilms, Frasassi Cave System, Italy. Appl Environ Microbiol 72: 5596–5609.

Macalady, J.L., Dattagupta, S., Schaperdoth, I., Jones, D.S., Druschel, G.K. and Eastman, D. (2008). Niche differentiation among sulfur-oxidizing bacterial populations in cave waters. ISME J 2: 590–601.

Markowitz, V.M., Chen, I.-M.A., Palaniappan, K., Chu, K., Szeto, E., Grechkin, Y., et al. (2012) IMG: the integrated microbial genomes database and comparative analysis system. Nucleic Acids Res 40: D115–D122.

Maugeri, T.L., Lentini, V., Spanò, A., and Gugliandolo, C. (2013) Abundance and Diversity of Picocyanobacteria in Shallow Hydrothermal Vents of Panarea Island (Italy). Geomicrobiology Journal 30: 93–99.

Mehta, M.P., Butterfield, D.A., and Baross, J.A. (2003) Phylogenetic Diversity of Nitrogenase (nifH) Genes in Deep-Sea and Hydrothermal Vent Environments of the Juan de Fuca Ridge. Appl Environ Microbiol 69: 960–970.

Meier, D.V., Pjevac, P., Bach, W., Hourdez, S., Girguis, P.R., Vidoudez, C., Aman, R. and Meyerdierks, A. (2017). Niche partitioning of diverse sulfur-oxidizing bacteria at hydrothermal vents. ISME J 11: 1545–1558.

Meyer, J.L. and Huber, J.A. (2013) Strain-level genomic variation in natural populations of Lebetimonas from an erupting deep-sea volcano. ISME J.

Mino, S., Kudo, H., Arai, T., Sawabe, T., Takai, K., and Nakagawa, S. (2014) Sulfurovum aggregans sp. nov., a hydrogen-oxidizing, thiosulfate-reducing chemolithoautotroph within the Epsilonproteobacteria isolated from a deep-sea hydrothermal vent chimney, and an emended description of the genus Sulfurovum. Int J Syst Evol Microbiol 64: 3195–3201.

Mintmier, B., McGarry, J.M., Sparacino-Watkins, C.E., Sallmen, J., Fischer-Schrader, K., Magalon, A., McCormick, J.R., Stolz, J.F., Schwarz, G., Bain, D.J. and Basu, P. Molecular cloning, expression and biochemical characterization of periplasmic nitrate reductase from *Campylobacter jejuni*, FEMS Microbiol Lett 365: fny151.

Miranda, P.J., McLain, N.K., Hatzenpichler, R., Orphan, V.J., and Dillon, J.G. (2016) Characterization of Chemosynthetic Microbial Mats Associated with Intertidal Hydrothermal Sulfur Vents in White Point, San Pedro, CA, USA. Front Microbiol 7:.

Mukhopadhyay, R. and Rosen, B.P. (2002) Arsenate reductases in prokaryotes and eukaryotes. Environ Health Perspect 110: 745–748.

Murdock, S., Johnson, H., Forget, N., and Juniper, S.K. (2010) Composition and diversity of microbial mats at shallow hydrothermal vents on Volcano 1, South Tonga Arc. Cah. Biol. Mar. 51: 407–413.

Mußmann, M., Hu, F.Z., Richter, M., de Beer, D., Preisler, A., Jørgensen, B.B., et al. (2007) Insights into the Genome of Large Sulfur Bacteria Revealed by Analysis of Single Filaments. PLoS Biol 5: e230.

Nakagawa, S. and Takai, K. (2008) Deep-sea vent chemoautotrophs: diversity, biochemistry and ecological significance. FEMS microbiology ecology 65: 1–14.

O’Brien, C.E., Giovannelli, D., Govenar, B., Luther, G.W., Lutz, R.A., Shank, T.M., and Vetriani, C. (2015) Microbial biofilms associated with fluid chemistry and megafaunal colonization at post-eruptive deep-sea hydrothermal vents. Deep Sea Research Part II: Topical Studies in Oceanography 121: 31–40.

Olivares, J., Bedmar, E.J., and Sanjuán, J. (2013) Biological Nitrogen Fixation in the Context of Global Change. MPMI 26: 486–494.

Parks, D.H., Imelfort, M., Skennerton, C.T., Hugenholtz, P., and Tyson, G.W. (2015) CheckM: assessing the quality of microbial genomes recovered from isolates, single cells, and metagenomes. Genome Res 25: 1043–1055.

Patwardhan, S., Foustoukos, D.I., Giovannelli, D., Yucel, M., and Vetriani, C. (2018) Ecological succession of sulfur-oxidizing Epsilon-and Gammaproteobacteria during colonization of a shallow-water gas vent. Front Microbiol 9: 2970. doi.org/10.3389/fmicb.2018.02970.

Pérez-Rodríguez, I., Bohnert, K.A., Cuebas, M., Keddis, R., and Vetriani, C. (2013) Detection and phylogenetic analysis of the membrane-bound nitrate reductase (Nar) in pure cultures and microbial communities from deep-sea hydrothermal vents. FEMS Microbiol Ecol 86: 256–267.

Pérez-Rodríguez, I., Bolognini, M., Ricci, J., Bini, E., and Vetriani, C. (2015) From deep-sea volcanoes to human pathogens: a conserved quorum-sensing signal in Epsilonproteobacteria. ISME J 9: 1222–1234.

Pitcher, R.S. and Watmough, N.J. (2004) The bacterial cytochrome cbb3 oxidases. Biochimica et Biophysica Acta (BBA) -Bioenergetics 1655: 388–399.

Pjevac, P., Meier D.V., Markert S., Hentschker C., Schweder T., Becher D., Gruber-Vodicka H.R., Richter M., Bach W., Amann R., Meyerdierks A. (2018) Metaproteogenomic Profiling of Microbial Communities Colonizing Actively Venting Hydrothermal Chimneys. Front Microbiol 9: 680. DOI=10.3389/fmicb.2018.00680.

Potter, L.C., Millington, P., Griffiths, L., Thomas, G.H., and Cole, J.A. (1999) Competition between Escherichia coli strains expressing either a periplasmic or a membrane-bound nitrate reductase: does Nap confer a selective advantage during nitrate-limited growth? Biochemical Journal 344: 77–84.

Price, R.E. and Giovannelli, D. (2017) A Review of the Geochemistry and Microbiology of Marine Shallow-Water Hydrothermal Vents. In, Reference Module in Earth Systems and Environmental Sciences. Elsevier.

Price, R.E., Lesniewski, R., Nitzsche, K., Meyerdierks, A., Saltikov, C., Pichler, T., and Amend, J. (2013) Archaeal and bacterial diversity in an arsenic-rich shallow-sea hydrothermal system undergoing phase separation. Front Microbiol 4:.

Quast, C., Pruesse, E., Yilmaz, P., Gerken, J., Schweer, T., Yarza, P., et al. (2013) The SILVA ribosomal RNA gene database project: improved data processing and web-based tools. Nucleic Acids Res 41: D590–D596.

Rathgeber, C., Yurkova, N., Stackebrandt, E., Beatty, J.T., and Yurkov, V. (2002) Isolation of Tellurite-and Selenite-Resistant Bacteria from Hydrothermal Vents of the Juan de Fuca Ridge in the Pacific Ocean. Appl Environ Microbiol 68: 4613–4622.

Rau, G.H. (1981) Low 15N/14N in hydrothermal vent animals: ecological implications. Nature 289: 484–485.

Reigstad, L.J., Jorgensen, S.L., Lauritzen, S.-E., Schleper, C., and Urich, T. (2011) sulfur-oxidizing Chemolithotrophic Proteobacteria Dominate the Microbiota in High Arctic Thermal Springs on Svalbard. Astrobiology 11: 665–678.

Reyes, C., Schneider, D., Lipka, M., Thürmer, A., Böttcher, M.E., and Friedrich, M.W. (2017) Nitrogen Metabolism Genes from Temperate Marine Sediments. Mar Biotechnol (NY) 19: 175–190.

Richardson, D.J., Berks, B.C., Russell, D.A., Spiro, S., and Taylor, C.J. (2001) Functional, biochemical and genetic diversity of prokaryotic nitrate reductases. CMLS, Cell Mol Life Sci 58: 165–178.

Ristova, P.P., Wenzhöfer, F., Ramette, A., Felden, J., and Boetius, A. (2015) Spatial scales of bacterial community diversity at cold seeps (Eastern Mediterranean Sea). The ISME Journal 9: 1306.

Rodriguez-R, L.M. and Konstantinidis, K.T. (2014) Bypassing Cultivation To Identify Bacterial Species: Culture-independent genomic approaches identify credibly distinct clusters, avoid cultivation bias, and provide true insights into microbial species. Microbe Magazine 9: 111–118.

Ruehland, C., Blazejak, A., Lott, C., Loy, A., Erséus, C., and Dubilier, N. (2008) Multiple bacterial symbionts in two species of co-occurring gutless oligochaete worms from Mediterranean sea grass sediments. Environ Microbiol 10: 3404–3416.

Schut, G.J., Bridger, S.L., and Adams, M.W.W. (2007) Insights into the Metabolism of Elemental Sulfur by the Hyperthermophilic Archaeon Pyrococcus furiosus: Characterization of a Coenzyme A-Dependent NAD(P)H Sulfur Oxidoreductase. J Bacteriol 189: 4431–4441.

Sharrar, A.M., Flood, B.E., Bailey, J.V., Jones, D.S., Biddanda, B.A., Ruberg, S.A., et al. (2017) Novel Large Sulfur Bacteria in the Metagenomes of Groundwater-Fed Chemosynthetic Microbial Mats in the Lake Huron Basin. Front Microbiol 8:.

Sievert, S. and Vetriani, C. (2012) Chemoautotrophy at Deep-Sea Vents: Past, Present, and Future. Oceanography 25: 218–233.

Sievert, S.M., Brinkhoff, T., Muyzer, G., Ziebis, W., and Kuever, J. (1999) Spatial heterogeneity of bacterial populations along an environmental gradient at a shallow submarine hydrothermal vent near Milos Island (Greece). Appl Environ Microbiol 65: 3834–3842.

Sievert, S.M., Ziebis, W., Kuever, J., and Sahm, K. (2000) Relative abundance of Archaea and Bacteria along a thermal gradient of a shallow-water hydrothermal vent quantified by rRNA slot-blot hybridization. Microbiology 146: 1287–1293.

Suess, E. (2014). Marine cold seeps and their manifestations: geological control, biogeochemical criteria and environmental conditions. Int J Earth Sci 103: 1889–1916.

Tarasov, V.G., Gebruk, A.V., Mironov, A.N., and Moskalev, L.I. (2005) Deep-sea and shallow-water hydrothermal vent communities: Two different phenomena? Chemical Geology 224: 5–39.

Veith, A., Botelho, H.M., Kindinger, F., Gomes, C.M., and Kletzin, A. (2012) The Sulfur Oxygenase Reductase from the Mesophilic Bacterium Halothiobacillus neapolitanus Is a Highly Active Thermozyme. J Bacteriol 194: 677–685.

Vetriani, C., Chew, Y.S., Miller, S.M., Yagi, J., Coombs, J., Lutz, R.A., and Barkay, T. (2005) Mercury Adaptation among Bacteria from a Deep-Sea Hydrothermal Vent. Appl Environ Microbiol 71: 220–226.

Vetriani, C., Voordeckers, J.W., Crespo-Medina, M., O’Brien, C.E., Giovannelli, D., and Lutz, R.A. (2014) Deep-sea hydrothermal vent Epsilonproteobacteria encode a conserved and widespread nitrate reduction pathway (Nap). ISME J.

Visser, J.M., de Jong, G.A.H., Robertson, L.A., and Kuenen, J.G. (1997) A novel membrane-bound flavocytochrome c sulfide dehydrogenase from the colourless sulfur bacterium Thiobacillus sp. W5. Arch Microbiol 167: 295–301.

Wagner, G.P., Kin, K., and Lynch, V.J. (2012) Measurement of mRNA abundance using RNA-seq data: RPKM measure is inconsistent among samples. Theory Biosci 131: 281–285.

Waite, D.W., Vanwonterghem, I., Rinke, C., Parks, D.H., Zhang, Y., Takai, K., et al. (2017) Comparative Genomic Analysis of the Class Epsilonproteobacteria and Proposed Reclassification to Epsilonbacteraeota (phyl. nov.). Front Microbiol 8:.

Wu, Y.-W., Tang, Y.-H., Tringe, S.G., Simmons, B.A., and Singer, S.W. (2014) MaxBin: an automated binning method to recover individual genomes from metagenomes using an expectation-maximization algorithm. Microbiome 2: 26.

Yamamoto, M. and Takai, K. (2011) Sulfur Metabolisms in Epsilon-and Gamma-Proteobacteria in Deep-Sea Hydrothermal Fields. Front Microbiol 2:.

Yamamoto Masahiro, Nakagawa Satoshi, Shimamura Shigeru, Takai Ken, and Horikoshi Koki (2010) Molecular characterization of inorganic sulfur-compound metabolism in the deep-sea epsilonproteobacterium Sulfurovum sp. NBC37-1. Environmental Microbiology 12: 1144–1153.

Yücel, M., Sievert, S., Vetriani, C., Foustoukos, D., Giovannelli, D. and Le Bris, N. (2013). Eco-geochemical dynamics of a shallow-water hydrothermal vent system at Milos Island, Aegean Sea (Eastern Mediterranean). Chem. Geol. 356:11–20.

